# A pan-cancer analysis of matrisome proteins reveals CTHRC1and LOXL2 as major ECM regulators across cancers

**DOI:** 10.1101/2021.04.27.441627

**Authors:** Keerthi Harikrishnan, Srinivas Seshagiri Prabhu, Nagaraj Balasubramanian

## Abstract

The extracellular matrix as part of the tumor microenvironment can regulate cancer cell growth and progression. Using TCGA data from 30 cancer types, the top 5% of matrisome genes with amplifications or deletions that affect survival in cancers were identified. Eight of these genes show altered expression in ~50% or more cancers affecting survival in ~20% or more. Among them SNED1 is the most downregulated and CTHRC1 and LOXL2 most upregulated. Differential gene expression analysis of SNED-1 did not identify any genes it regulates across cancers, while CTHRC1 and LOXL2 affected 19 and 5 genes respectively in 3 or more cancers. STRING analysis of these genes classified them as ‘extracellular’, involved prominently in ECM organization. Their correlation and co-occurrence in context of their effect on survival and staging of the disease identified MMP13, POSTN and SFRP4 along with COL11A1, COL10A1, COL1A1, ADAMTS12 and PPAPDC1A as possible interactors of CTHRC1 and LOXL2 in cancers. These are implicated in collagen organization, making it vital to matrisome regulation of cancers. Clinical Proteomic Tumor Analysis Consortium data confirms the changes in expression of these genes along with CTHRC1 and LOXL2 in breast and lung cancer, further supporting their implication as vital pan-cancer matrisome mediators.

**Highlights:**

- CTHRC1 and LOXL2 are prominently upregulated pan-cancer matrisome genes.
- High CTHRC1 and LOXL2 expression is associated with disease progression and poor survival in cancers.
- CTHRC1 with POSTN, MMP13 and SFRP4 and LOXL2 with COL11A1, COL10A1, COL1A1, ADAMTS12 and PPAPDC1A drive matrisome regulation of cancers.
- CTHRC1 and LOXL2 could prominently drive collagen organization and function across cancers.

## Introduction

Extracellular matrix (ECM) is a dynamic interconnected mesh of macromolecules which provides structural support and also regulates cellular behavior via mechanical and biochemical cues. It regulates several cellular processes including proliferation, differentiation, migration, invasion and survival [1]. ECM composition is tightly regulated by the cells and changes in ECM production, secretion, deposition and remodeling are reported in pathological diseases such as atherosclerosis, fibrosis, skeletal disorders, vascular disorders and cancer [2,3]. ECM composition also varies between the tumor cells, tumor stroma and it is distinctly different across the metastatic sites [4,5]. Recent studies characterizing changes in the composition of ECM in normal and tumor microenvironments have emphasized the importance of ECM and its contribution towards developing novel biomarkers and therapeutic targets [5,6][7].

The “Matrisome” pioneered by *Naba et al*., is an ensemble of genes that codes for core ECM proteins, ECM associated growth factors, ECM regulators and other ECM associated factors [6,8]. It accounts for 4% of human and mouse genome and reflects the composition of proteins in normal and tumor tissues [3,9]. Since the publication of MatrisomeDB in 2012, the understanding of the role of ECM in cancers has significantly enhanced. Matrisome proteins like IGFBP3, IGFBP4, IGFBP5, CCN family, THBS2, TNN and VWA9 are detected primarily in cancer tissues [2] while LOXL2 [10], COMP, POSTN[11], TNC[12], TNX[12] and FN (EIIIA and EIIIB variant) [13] have been reported to be upregulated in cancers. Oncomine analysis of core matrisome genes in lung, gastric, ovarian and colon cancers shows that a signature of 9 genes (COL11A1, SPP1, MFAP2, COL10A1, BGN, COMP, AGRN and MXRA5) is associated with poor survival and is involved in regulating cancer hallmarks such as epithelial to mesenchymal transition (EMT), and angiogenesis [14]. Data mining of 10 NSCLC microarray datasets has identified 29 ECM signature genes which were found to be consistently upregulated in patients with NSCLC and also predicts prognosis [15]. Analysis of 12 cancer types (lung, pancreas, prostate, kidney, stomach, colon, ovary, breast, liver, bladder and skin) shows that tumor matrisome index (TMI) is associated with disease progression and poor clinical outcome [16]. In addition, tumors with high TMI show an enrichment for MAGEA3 and CD8 positive T cells and also display high expression of B7-H3 which is negatively associated with clinical outcome in solid tumors [17].

Pan-Cancer analysis of TGF ß associated ECM gene expression shows a set of matrisome genes to be upregulated in cancer and the expression is associated with worse prognosis. This study also reveals an association of aberrant ECM expression with immunosuppression in cancers [16,18]. Cell-cell adhesion, FOXO, Wnt pathways are found to control matrisome in most cancer types whereas TP53, Notch and TGFß signaling pathways regulate matrisome genes in some cancers [19].

Using a multiomics approach and machine learning, several landmark matrisome genes have been identified from 74 clinical and molecular subtypes of cancers that show prognostic significance [20]. Bioinformatic analysis of the copy number alterations (CNA’s) reveals that matrisome genes display a disproportionately high number of CNA’s and mutations compared to the rest of the genome [21] across cancers. This increase in the genome alterations of matrisome was further predictive of prognosis across cancer types. Together, these findings highlight a significant role for ECM genes in cancer progression.

While recent studies have evaluated the role of matrix proteins as a group or ECM index, our study was aimed at identifying individual matrisome genes that act as vital ECM regulators in the tumor microenvironment. A pan-cancer analysis of copy number variation, expression, survival of matrisome proteins has identified CTHRC1 and LOXL2 as major pan-cancer ECM regulators. These studies further reveal novel candidate genes POSTN, MMP13, SFRP4 and COL11A1, COL10A1, COL1A1, ADAMTS12, PPAPDC1A to possibly work with CTHRC1 and LOXL2 in regulating cancers.

## Materials and Methods

### Data sources

The list of matrisome genes was downloaded from the matrisome database (http://matrixdb.univ-lyon1.fr/). TCGA Pan-Cancer copy number data calculated using the GISTIC2 threshold method was downloaded from the UCSC Xena browser (https://xenabrowser.net/). TCGA and GTEx data was used to perform the expression, survival, correlation, cooccurrence disease stage and protein analysis. The analysis was restricted to 30 types of cancers from TCGA which include: Adrenocortical Carcinoma (ACC), Bladder Urothelial Carcinoma (BLCA), Breast Invasive Carcinoma (BRCA), Cervial Squamous Cell Carcinoma (CESC), Cholangiocarcinoma (CHOL), Colon Adenocarcinoma (COAD), Lymphoid Neoplasm Diffuse Large B-cell Lymphoma (DLBC), Esophageal Carcinoma (ESCA), Glioblastoma Multiforme (GBM), Head and Neck Squamous Cell Carcinoma (HNSC), Kidney Chromophobe (KICH), Kidney Renal Clear Cell Carcinoma (KIRC), Kidney Renal Papillary Cell Carcinoma (KIRP), Brain Lower Grade Glioma (LGG), Liver Hepatocellular Carcinoma (LIHC), Lung Adenocarcinoma (LUAD), Lung Squamous Cell Carcinoma (LUSC), Ovarian Serous Cystadenocarcinoma (OV), Pancreatic Adenocarcioma (PAAD), Pheochromocytoma and Paraganglioma (PCPG), Prostate Adenocarcinoma (PRAD), Rectum Adenocarcinoma (READ), Sarcoma (SARC), Skin Cutaneous Melanoma (SKCM), Stomach Adenocarcinoma (STAD), Testicular Germ Cell Tumors (TGCT), Thyroid Carcinoma (THCA), Thymoma (THYM), and Uterine Corpus Endometrial Carcinoma (UCEC), Uterine Carcinosarcoma (UCS).

#### Copy number variation Analysis

First genes were clustered based on their functionality (ECM genes, Proteoglycans etc.). These gene clusters were prepared as an excel file containing a single column with gene symbols as cell entries and this list represented the genes of interest whose copy number variations were to be analyzed. There were 10845 samples in total and the gene-level copy number estimate values of −2, −1, 0, 1, 2 represented deep deletion, shallow deletion, no change, amplification and gain respectively. The excel file retrieved after extracting the dataset was in the genomic Matrix format (ROWs (identifiers) x COLUMNs (samples)). Next, we wrote a code in Python to find the gene-level copy number estimate values of our genes of interest across all 10845 samples. To achieve this, we had to explore our list in the database and retrieve the corresponding values for the samples and process the data to calculate the number of deep deletion and gain. The top 5% of the genes (n=104) with amplification or deep deletion were then used for further analysis. The code used to perform the analysis can be made available upon written request.

### Expression, Survival, Correlation and Disease stage Analysis

RNA expression for the matrisome genes was analyzed using GEPIA2 portal (http://gepia2.cancer-pku.cn/#analysis) which contains the expression data for 9736 tumor samples and 8587 normal samples and the data is processed using a standard processing pipeline. The data from 30 cancer types is represented in a box plot (log scale) as mean ± standard deviation (S.D). We also evaluated the expression of the matrisome genes by pathological stage using the stage plot function in the database. The data from 30 cancer types is represented in violin plot (log scale) which shows the distribution of data. p value less than 0.05 was considered statistically significant. Survival analysis for all the matrisome genes was performed using GEPIA2 portal using a custom cutoff (High 75 percentile, Low 25 percentile). Log rank p values and the hazard ratio (HR) with 95% confidence interval was calculated using the web portal and a p value of less than 0.05 was considered statistically significant. Percentage alteration based on expression and survival data was calculated as follows:

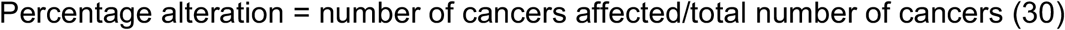

Survival analysis for Figure 2B, 3B was performed using the survival analysis (Analysis 2, https://GitHub.com/mdozmorov/TCGAsurvival/blob/master/survival.Rmd) code in the TCGA2STAT package in R (version 3.6.3). Correlation analysis for CTHRC1 along with its 10 hub genes across 30 cancer types was done using the spearman correlation coefficient. p value of less than 0.05 was considered statistically significant.

### Mutation and Cooccurrence Analysis

Mutation analysis for 8 shortlisted genes (based on expression, survival data as mentioned above) was analyzed using the cBioPortal database across 30 cancer types. It showed the mutational burden along with the copy number analysis in the patient samples. We also used this portal to identify the co-expressed genes of CTHRC1 and LOXL2 (30 cancer types) in the cBioPortal using the cooccurrence-mutual exclusivity tab in the database. p less than or equal to 0.05 was considered statistically significant.

### Differential Gene Expression Analysis

First the data was preprocessed locally in R using the preprocessing script. The differential gene expression (DEG) analysis was performed (30 cancer types) in groups with high expression (75%) or low (25%) expression. Threshold for DEG’s were set at fold change > 2, −2 with a p value of 0.05. The following source code from the GitHub repository was used to perform the preprocessing and the DEG analysis. (https://GitHub.com/mdozmorov/TCGAsurvival/blob/master/misc/TCGA_preprocessing.R, https://GitHub.com/mdozmorov/TCGAsurvival/blob/master/TCGA_DEGs.Rmd)

#### Venn Diagram Analysis for overlapping genes

Omics Box software and the Venn diagram tool (http://bioinformatics.psb.ugent.be/webtools/Venn/) were used to identify the overlapping differentially expressed genes across cancer types. 20 genes that overlapped in 3 or more cancers were used for further analysis. Venn diagram tool was also used to identify the top hub genes that overlapped in survival, disease stage expression, correlation and cooccurrence analysis. 3 genes identified were then used to perform CPTAC analysis.

#### Protein protein interaction and functional enrichment analysis

STRING database (https://string-db.org/) was used to identify the protein protein interactions of CTHRC1, LOXL2 and SNED1 along with the differentially expressed genes across different cancer types. The list of proteins along with CTHRC1, LOXL2 and SNED1 was entered in the online database and a medium confidence interaction score of 0.4 was used to generate the full string network. The network was then exported as a high-resolution image. Functional enrichment analysis performed using the STRING database included Biological process, molecular function, cellular compartment and KEGG pathways. FDR of < 0.05 was used to identify the gene ontology (GO) terms that were statistically significant.

### CytoHubba Analysis

The PPI network constructed using the STRING database was sent to Cytoscape using the web link. Cytoscape (version 3.8.2) is an open-source software used for visualization and analysis of protein protein interaction networks. Using the cytoHubba tool in the software, we identified the top 10 hub genes base on the degree of connectivity, betweenness centrality, stress, closeness centrality, and clustering coefficient. These 10 hub genes were then used for further analysis.

### UACLAN analysis

UACLAN is a web-based tool for analysis of CPTAC data from the TCGA cancer types. We used UACLAN database (http://manualcan.path.uab.edu/index.html) to analyze the protein levels of CTHRC1, POSTN, MMP13, SFRP4, LOXL2, COL11A1, COL10A1, COL1A1, ADAMTS12, and PPAPDC1A in breast and lung cancers. Data was represented as mean ± S.D and a p value of less than 0.05 was considered statistically significant.

## Results

### Pan-cancer analysis identifies top 8 altered matrisome genes

In this study, we first analyzed the copy number variation of matrisome genes (n= 1027) using the Pan-Cancer dataset from the UCSC cancer genome browser (***Figure 1A***). Overall, there were about 10845 samples in the dataset which were scored based on their copy number variations. We were interested in genes that had significant deep deletions or amplification (***Figure 1A***) and calculated their score for each gene. The top 5% of genes with deep deletions (>500) or amplification (>3800) were shortlisted to identify 104 genes. Using GEPIA2 portal, we evaluated the mRNA expression of these 104 genes and their effect on survival across 30 cancer types (***Figure 1A,1B and 1C***). For ease in data interpretation, gene expression was represented in three categories: upregulated, downregulated or no change in expression (***Figure 1B***). Similarly, the survival data was categorized as “yes” (gene expression associated with significant effect on survival), “no” (gene expression not associated with significant survival) and not known (***Figure 1C***). We then shortlisted genes whose expression was altered in 50% or more of cancers (n=17) and which affected survival in 20% or more of cancers (n=24; ***Figure 1D, E***). When overlapped, this identified 8 genes, CTHRC1, LOXL2, SNED1, IGFBP3, BMP1, PI3, COL1A2 and AEBP1 (***Figure 1D, E***). Copy number variation and mutation analysis for the 8 genes in cBioPortal showed CTHRC1 to be prominently amplified and SNED1 to be prominently deleted across cancers (***Figure 1G***). These 8 genes were ranked based on relative expression and effect on survival (***Figure 1F***) and show SNED1 and CTHRC1, LOXL2 to be the top matrisome genes downregulated and upregulated respectively in cancers.

**Figure 1.**
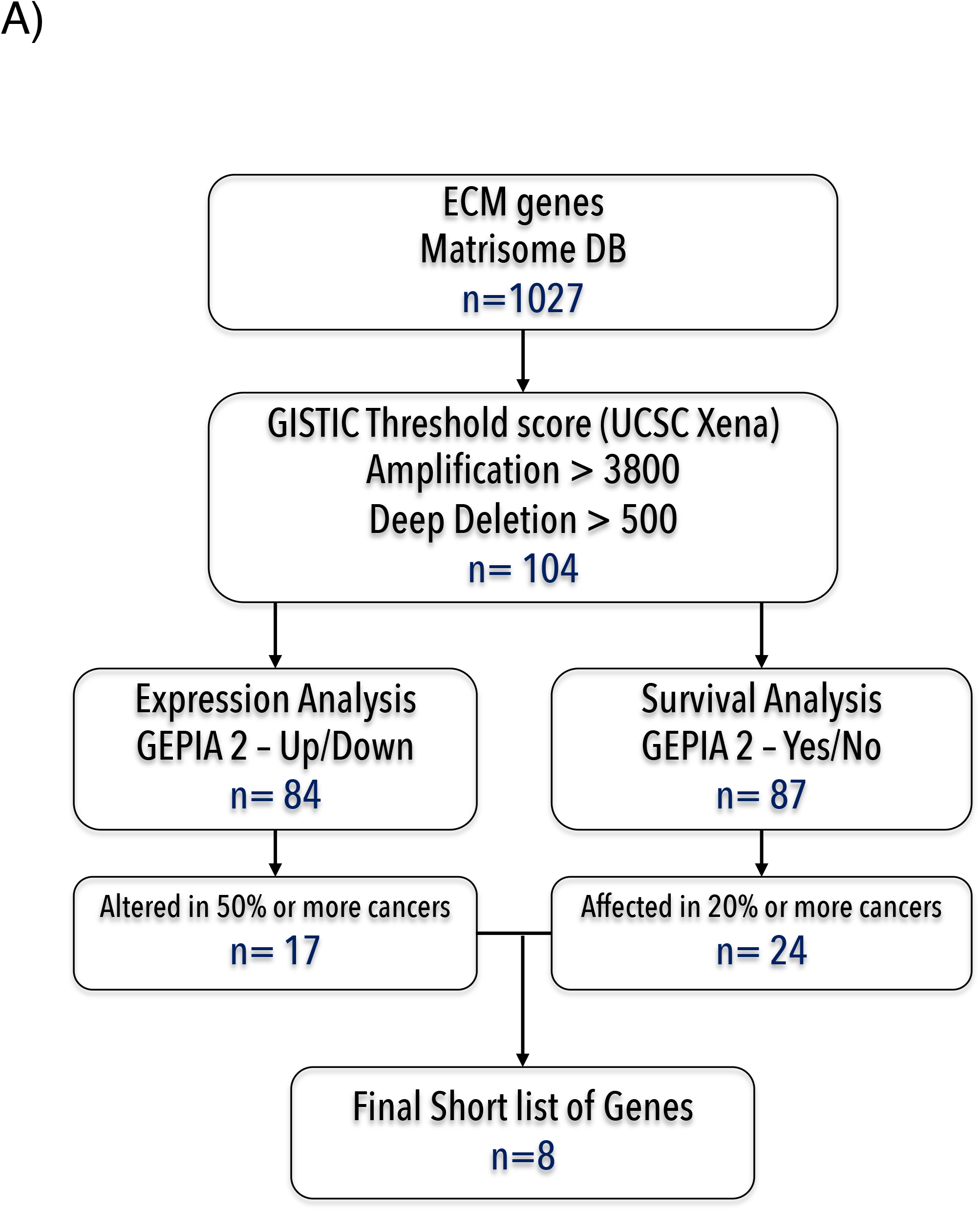
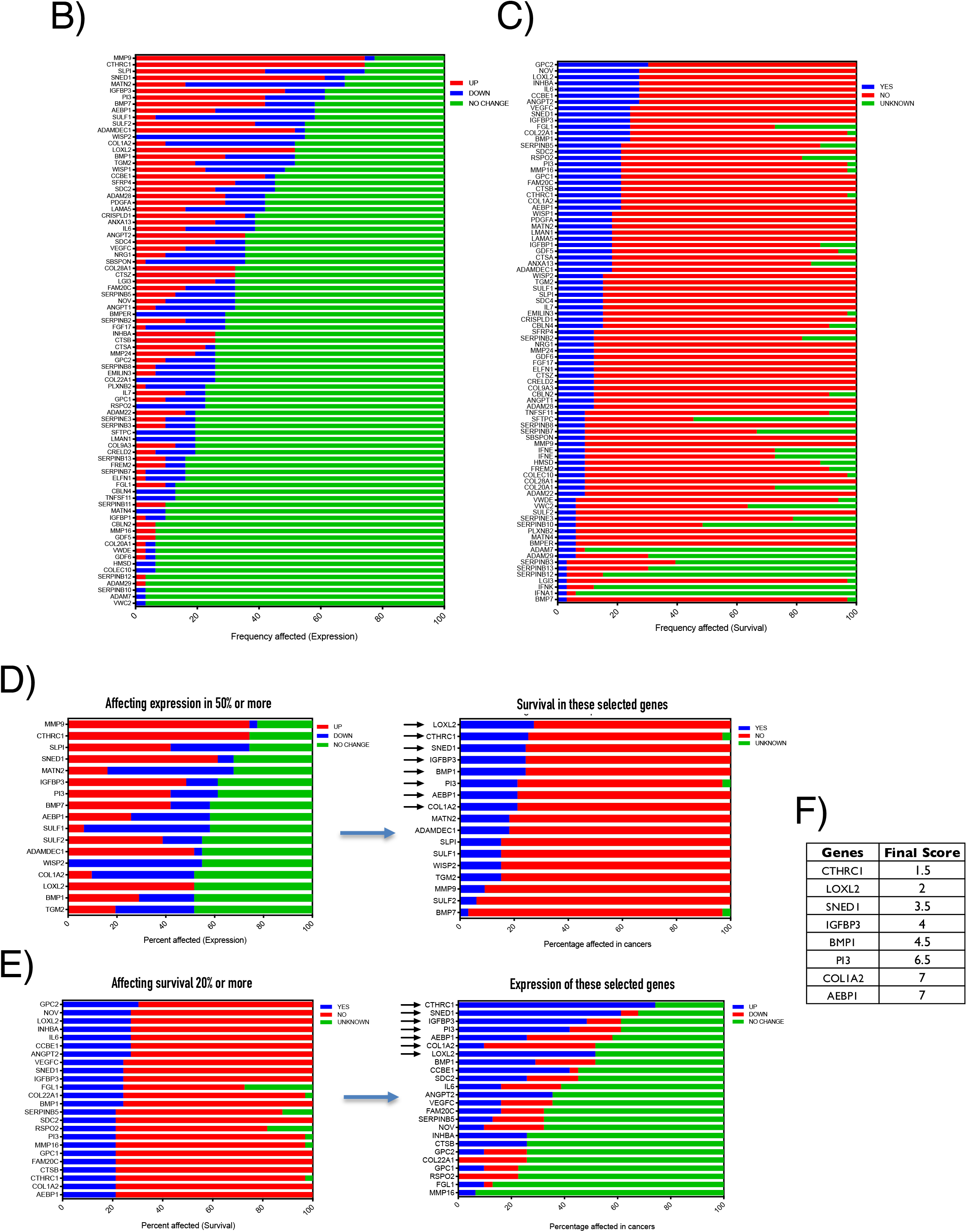
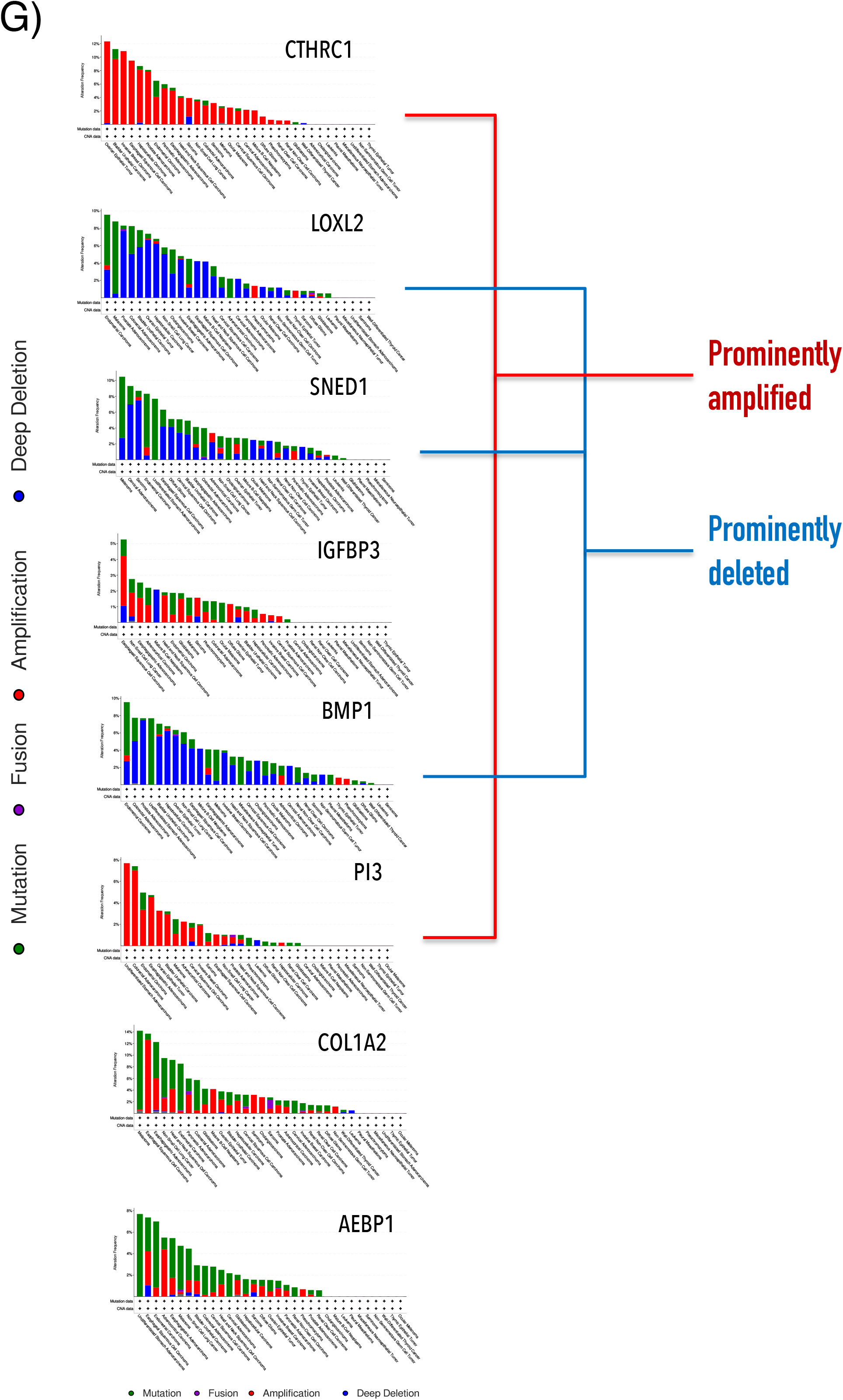
Expression, survival and mutation analysis of top matrisome genes in pan-cancers. **A)** Schematic shows the steps involved in shortlisting genes based on copy number, expression and survival in the study. **B)** Bar graph shows the frequency of altered gene expression for the top 5% of matrisome genes arranged in the order of high to low. (RED – upregulated in cancers, GREEN - downregulated in cancers, BLUE – no change in expression. **C)** Bar graph shows the frequency of genes affecting survival for the top 5% of matrisome genes arranged in the order of high to low. (BLUE – significant effect on survival, RED – no significant effect on survival, GREEN – not known). **D)** Bar Graph shows the expression of 17 genes whose expression is altered in 50% or more cancers (BLUE - Downregulated, RED - upregulated, GREEN - no change) (left panel), looking at their impact on survival (BLUE - significant effect on survival, RED – no significant effect on survival, GREEN – not known) (right panel). They are arranged in the arranged in the order of high to low. Arrows identify genes whose expression is altered in ≥50% and also affects survival in ≥ 20% of cancers. **E)** Bar Graph shows the listing of 24 genes that affect survival in 20% or more cancers (BLUE - significant effect on survival, RED – no significant effect on survival, GREEN – not known) (left panel), looking at the expression of these genes (BLUE - Downregulated, RED-upregulated, GREEN-no change) (left panel) (right panel). Arrows identify genes whose expression is altered in ≥50% and also affects survival in ≥ 20% of cancers. **F)** Listing of 8 genes with their score calculated as described in methods based on their position in the expression (**D**) and survival (**E**) graphs. **G)** Mutational and copy number alterations of 8 genes (CTHRC1, LOXL2, SNED1, IGFBP3, BMP1, PI3, COL1A2 and AEBP1) in 30 cancer types. The alterations include mutations (GREEN), fusion (PURPLE), amplification (RED) and deep deletions (BLUE).

### Pan-cancer analysis of SNED1

SNED1(Sushi, nidogen and EGF like domain containing protein-1) is a glycoprotein known to be involved in embryonic development [22]. Our pan-cancer analysis of matrisome genes differentially expressed and affecting survival in TCGA cancer types, identified SNED1 to be significantly downregulated in bladder, breast, colon, brain, kidney, lung, ovarian, adrenal, prostate, rectal, skin, testicular and uterine cancers while it was upregulated in only thymus cancer (***Figure 2A, C***). We further evaluated the effect of SNED1 on survival using the TCGA2STAT package in R studio which showed low levels of SNED1 to be associated with poor survival in bladder, cervical, esophageal, brain, kidney, liver, ovarian, adrenal, thymus and uterine cancers (***Figure 2B, C***). Differential gene expression analysis in bladder, brain (GBM, LGG), ovarian and adrenal cancers and selection for the top 5% genes with a fold change greater or lesser than 2 (p<0.05) was done to perform further downstream analysis (***Figure 2C***). Screening of these genes for their potential overlap across these cancers revealed only HSPB7 to be the common downstream mediator in bladder and adrenal cancers. No common mediator was detected in 3 or more cancers that we were using as a criterion to implicate them strongly across cancers (***Figure 2D***).

**Figure 2.**
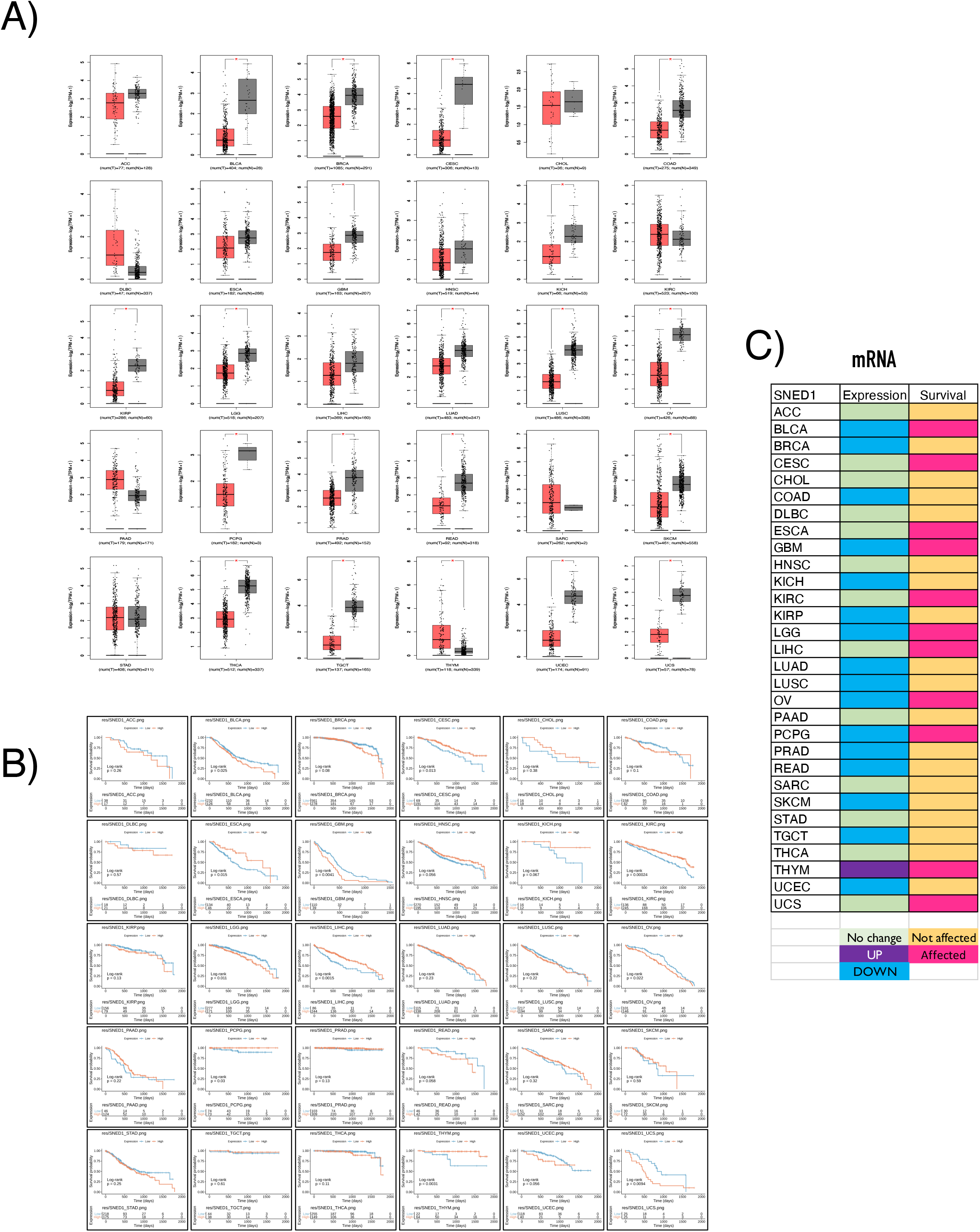
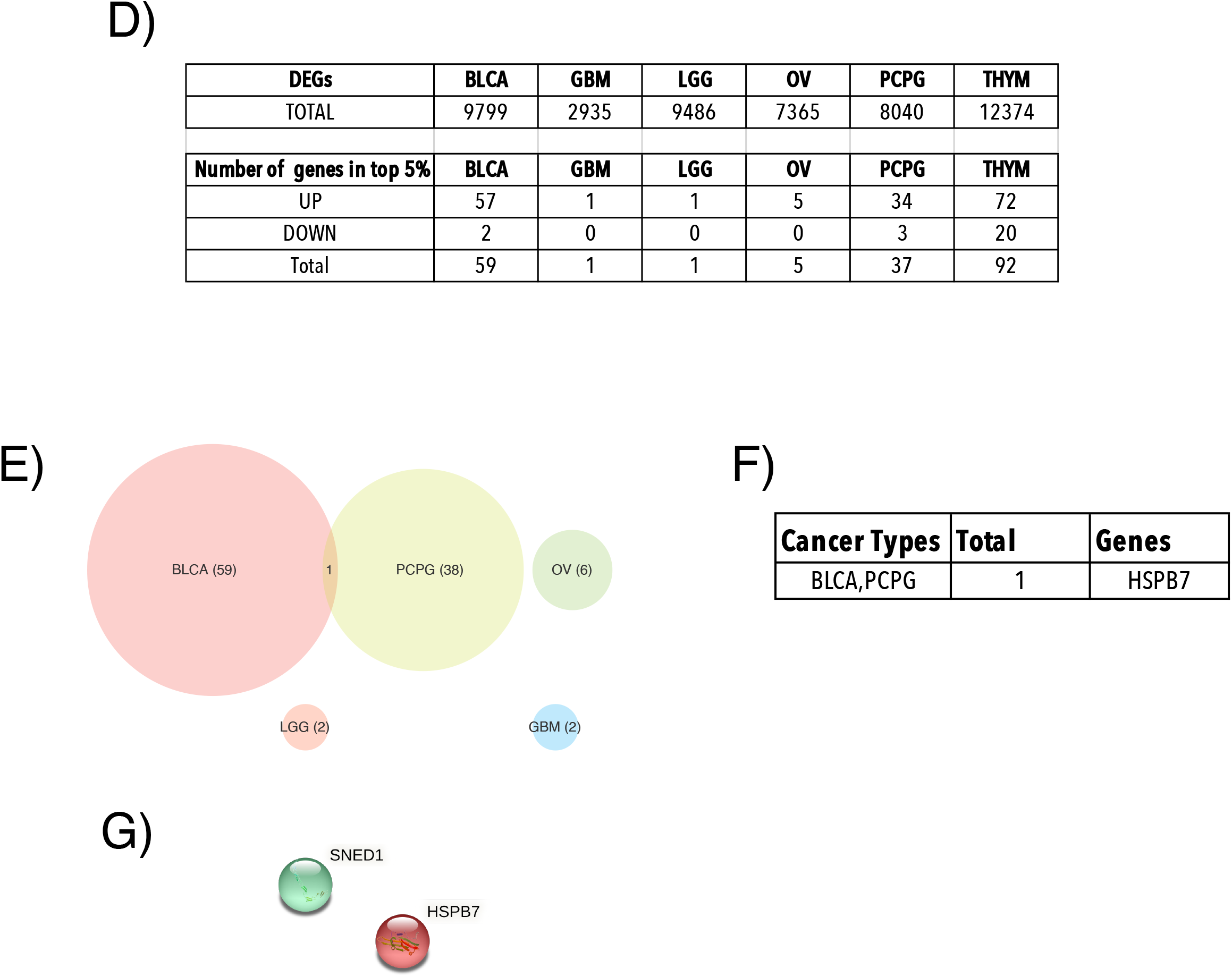
Pan-Cancer analysis of SNED1. **A)** Graphs show expression of SNED1 in 30 different cancer types from GEPIA2 database (GREY – normal, RED – tumor). Expression data is represented as mean ± standard deviation (S.D) on a log scale using a box plot. p value of < 0.05 was determined as statistically significant. **B)** Graphs show effect of SNED1 on survival in 30 different cancer types using R studio (BLUE – low expression, RED – high expression). p value of < 0.05 was determined as statistically significant. **C)** Table shows SNED1 expression and survival data in 30 cancers. Upregulated expression is marked in PURPLE, downregulated expression in BLUE and No change in expression marked in GREEN. Effect seen on survival is marked in PINK and lack of effect marked in CANTALOUPE. **D)** Table lists the TOTAL number of Differential expressed Genes (DEG) in 5 selected cancers (BLCA, GBM, LGG, OV, PCPG – as detailed in Methods). The lower table lists the number of differentially expressed genes in the top 5% with a fold change of >2 (as detailed in methods) that are up regulated (UP) or downregulated (DOWN). Their aggregate numbers are listed as TOTAL. **E)** This Venn diagram shows the overlap of these differentially expressed genes in the above listed 5 cancers. **F)** Table lists the only overlapping gene between multiple cancers as HSPB7. **G)** Protein protein interaction network constructed for SNED1 and HSPB7 using the STRING database shows no interaction between them.

### Pan-cancer analysis of LOXL2

Lysyl oxidase like 2 (LOXL2) belongs to the LOX family of proteins which are known to be involved in covalent crosslinking of elastin and collagen to stabilize ECM network [23]. Our analysis revealed that LOXL2 is upregulated in cancers (***Figure 1F***). LOXL2 expression evaluated using GEPIA2 portal confirmed its overexpression in bile duct, lymphoid, esophagus, brain, head & neck, kidney, liver, pancreas, adrenal, stomach, skin, testicular, thyroid and uterine cancers (***Figure 3A, 3C)***. Survival analysis showed high LOXL2 levels to be associated with poor survival in adrenal, bladder, breast, cervical, brain, head & neck, kidney, liver, lung, pancreas, stomach and thyroid cancers (***Figure 3B,3C)***. DEG analysis of brain, head & neck, kidney, liver, pancreas and stomach cancers was done and the top 5% genes with a fold change greater or lesser than 2 (p<0.05) selected for further analysis (***Figure 3D***). This identified 5 genes that were upregulated in 3 or more cancers (***Figure 3E***), with no genes seen to be downregulated. The 5 upregulated genes include COL11A1, COL10A1, COL1A1, ADAMTS12 and PPAPDC1A (***Figure 3E***). They were used to construct a protein protein interaction network using the STRING database (***Figure 3F)***. Functional enrichment analysis of this network revealed collagen fibril organization and ECM organization to be among the top 5 biological processes regulated by these network genes (***Figure 3G)***. Proteins in this network prominently belonged to the ECM compartment (***Figure 3H**).* KEGG analysis of this network further shows their involvement in protein digestion pathway (***Figure 3H)***. Since our DEG analysis identified only 5 genes to be differentially regulated by LOXL2 in 3 or more cancers, we hence decided to evaluate all the 5 genes for downstream analysis. To validate the significance of LOXL2 and the differentially regulated genes, we first performed a pan-cancer survival analysis of the 5 genes along with LOXL2 using GEPIA2. Of these genes, overexpression of COL11A1, COL10A1, COL1A1, ADAMTS12 and PPAPDC1, like LOXL2, is associated with poor prognosis (***Figure 3I**)*. We used GEPIA2 to determine if the change in expression of these genes and LOXL2 is associated with tumor grade. Statistical analysis of this change by one way ANOVA showed all 5 genes and LOXL2 expression to indeed be tumor stage dependent (***Figure 3J**)*. Spearman correlation analysis showed a positive correlation between these 5 genes and LOXL2 (***Figure 3K**)*. cBiportal analysis also indicated a co-occurrence between LOXL2 and COL11A1, COL10A1, COL1A1, ADAMTS12 and PPAPDC1A (***Figure 3L**)*. Combining the survival, tumor grade, co-occurrence and correlation analysis, we identified these 5 genes to be the most likely mediators of LOXL2 dependent function in cancers (***Figure 3M**)*.

**Figure 3.**
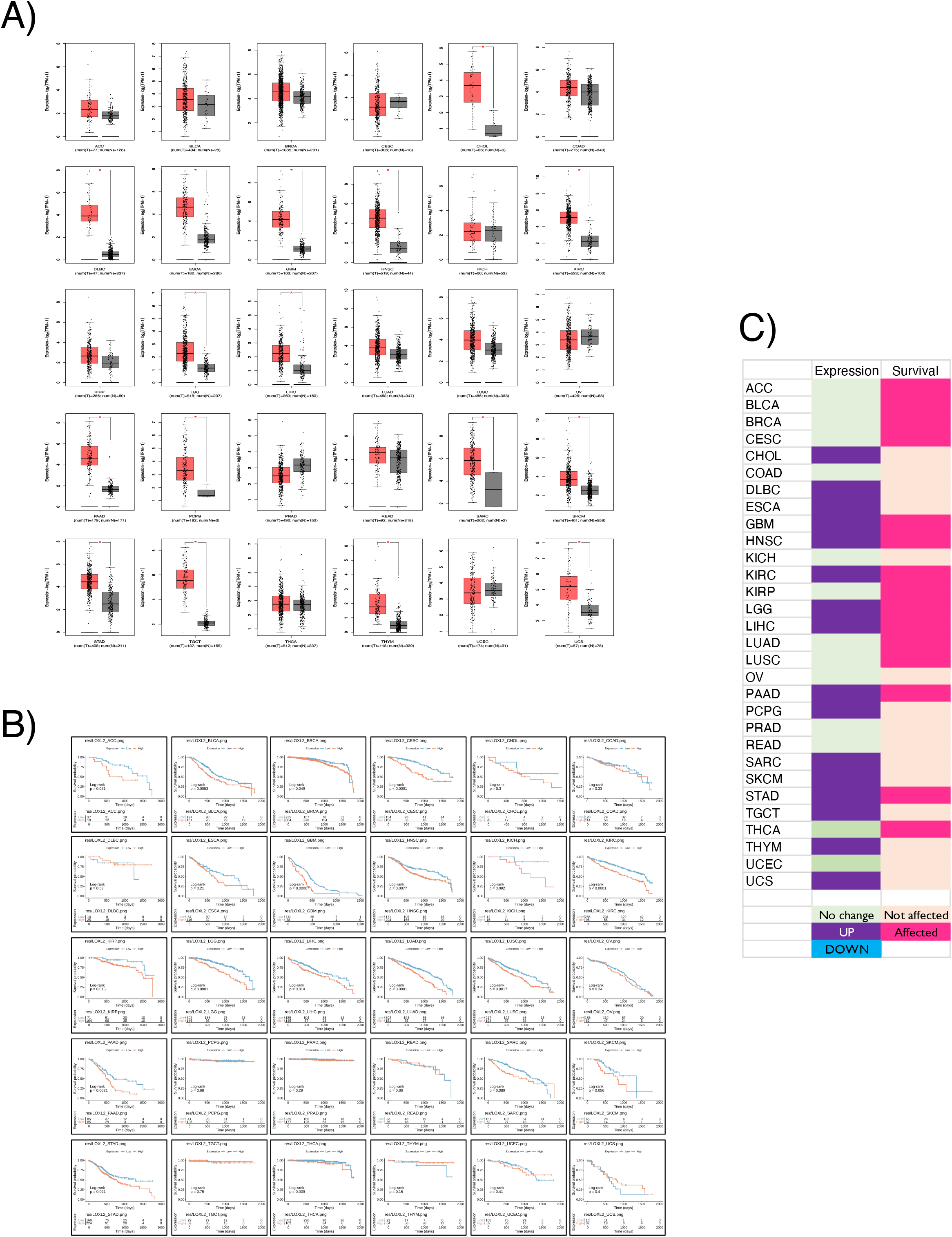
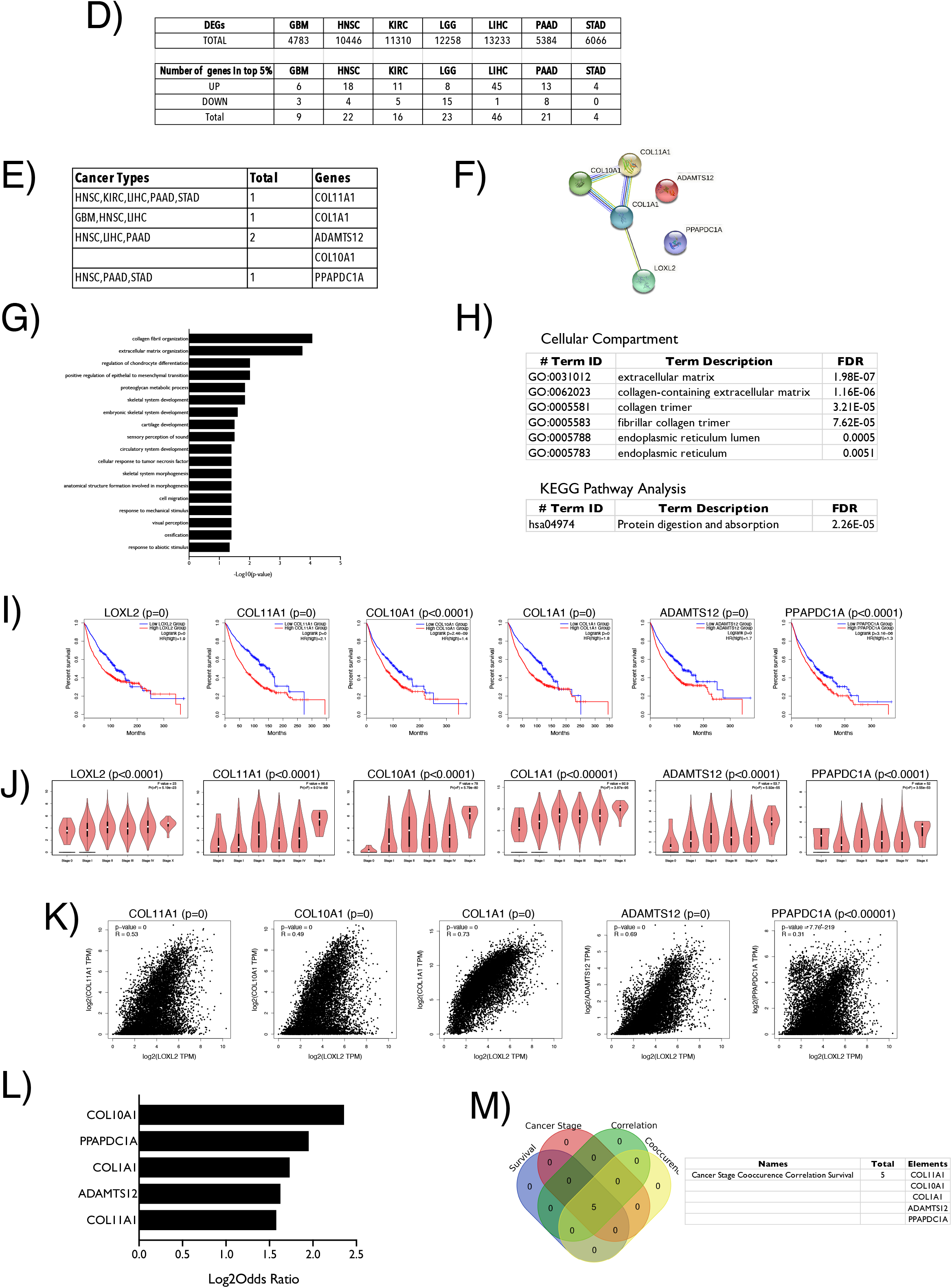
Pan-Cancer analysis of LOXL2 and validation of its differentially expressed genes. **A)** Graphs show expression of LOXL2 in 30 different cancer types from GEPIA2 database (GREY – normal, RED – tumor). Expression data is represented as mean ± standard deviation (S.D) on a log scale using a box plot. p value of < 0.05 was determined as statistically significant. **B)** Graphs show effect of LOXL2 on survival in 30 different cancer types using R studio. (BLUE – low expression, RED – high expression). p value of < 0.05 was determined as statistically significant. **C)** Table shows LOXL2 expression and survival data in 30 cancers. Upregulated expression is marked in PURPLE, downregulated expression in BLUE and No change in expression marked in GREEN. Effect seen on survival is marked in PINK and lack of effect marked in CANTALOUPE. **D)** Table lists the TOTAL number of Differential expressed Genes (DEG) in 7 selected cancers (GBM, HNSC, KIRC, LGG, LIHC, PAAD, STAD – as detailed in Methods). The lower table lists the number of differentially expressed genes in the top 5% with a fold change of >2 (as detailed in methods) that are up regulated (UP) or downregulated (DOWN). Their aggregate numbers are listed as TOTAL. **E)** Table lists the genes identified to be upregulated in 3 or more cancers. Cancer types are as listed and detailed in methods section. **F)** Protein protein interaction network constructed for these 5 differentially expressed genes with LOXL2 using the STRING database shows interactions between LOXL2 and COL1A1 and also among the differentially expressed genes. BLUE line marks predicted interactions from gene cooccurrence data, GREEN line marks predicted interactions based on gene neighborhood evidence, PURPLE line marks experimentally determined known interactions, YELLOW line marks interactions based on text mining and the LIGHT BLUE line marks interactions based on database evidence. **G)** Gene ontology terms enriched for biological process from the above STRING network analysis are listed in their descending order of significance. **H)** Table lists the gene ontology terms enriched for cellular compartments. Lower table lists the gene ontology terms enriched for protein digestion and absorption using KEGG pathway analysis. The False Discovery Rate (FDR) values from this analysis are listed for each gene ontology term. **I)** Graph represents percentage survival of LOXL2 and its 5 differentially expressed genes (COL11A1, COL10A1, COL1A1, ADAMTS12 and PPAPDC1A) in 30 cancers using GEPIA2 database. Graph in Blue and Red show the percentage survival over time for cancers with low and high expression respectively of the gene of interest. p values are as indicated above the graph. p values = 0 reflects very high significance. **J)** Violin plot shows the expression of LOXL2 and each of its 5 interactors across pathological stages in 30 cancers analyzed using GEPIA2. Significance comparing the stages of cancer for each gene of interest was calculated by ANOVA and significance reported. p values are as indicated above the graph. **K)** Scatter plots show the spearman correlation analysis for LOXL2 and its interactors in 30 cancers using GEPIA2. p values are as indicated above the graph. p values = 0 are representative of very high significance. **L)** Bar graphs shows log2 odds ratio from cBioPortal for LOXL2 determining its cooccurrence or mutual exclusivity with the 5 identified genes of interest (COL11A1, COL10A1, COL1A1, ADAMTS12 and PPAPDC1A) in 30 cancer types. **M)** This Venn diagram shows the overlap between genes that in a statistically significant manner affect survival, cancer staging, are related in correlation analysis and show cooccurrence in 30 cancer types. Table lists the 5 genes overlapping in all four analysis.

A recent study shows matrisome proteins in breast and lung cancers to display more copy number alterations compared to other cancers [21]. We hence evaluated the UACLAN portal data for breast and lung cancer and find LOXL2 protein levels to be upregulated in both cancers (***Figure 4A)***. In breast cancer samples LOXL2 was overexpressed with COL11A1, COL10A1 and ADAMTS12 (***Figure 4B**)*, while in lung cancers COL11A1, COL10A1, COL1A1 and ADAMTS12 were overexpressed relative to normal tissues (***Figure 4C**)*.

**Figure 4.**
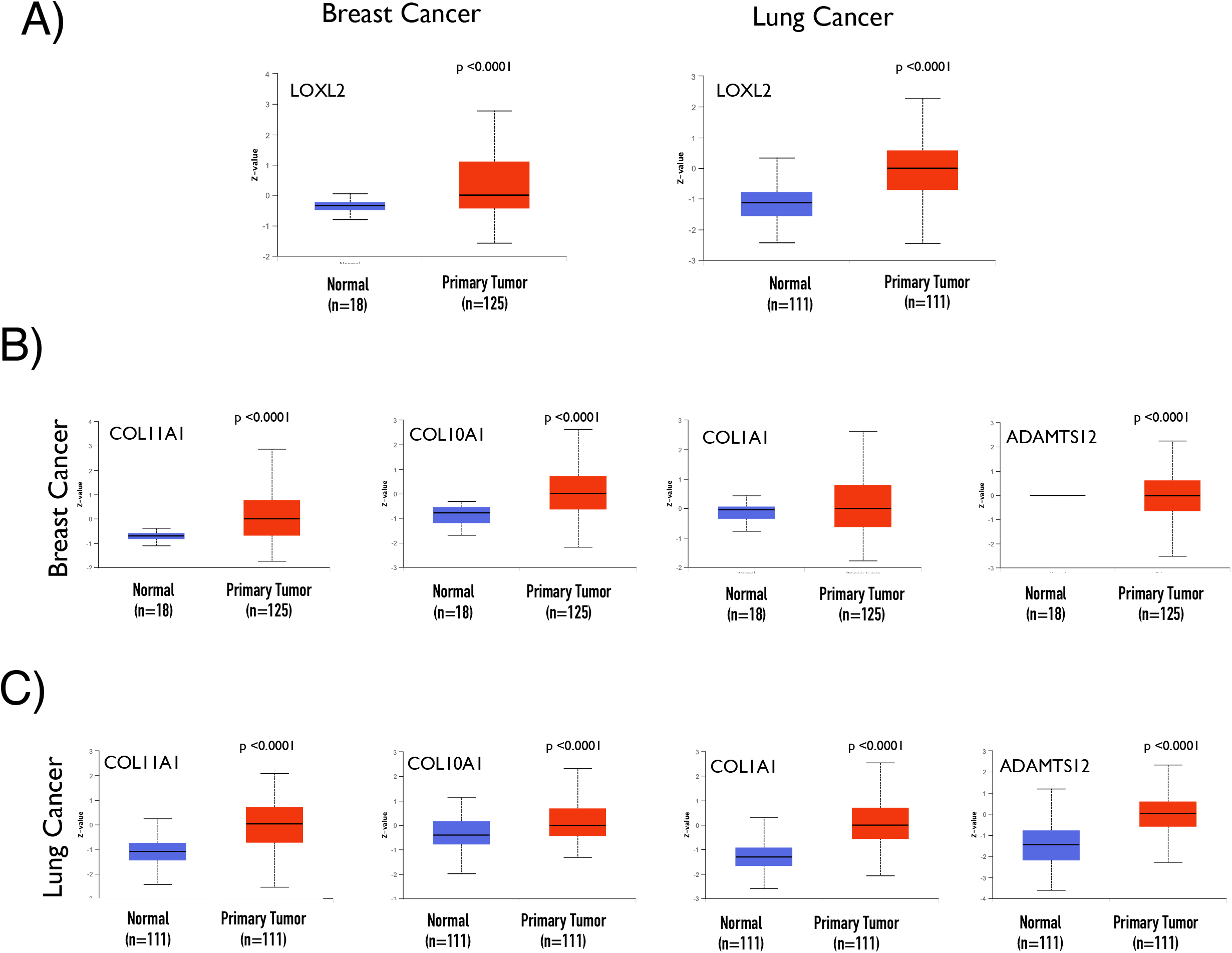
Protein levels of LOXL2, COL11A1, COL10A1, COL1A1 and ADAMTS12 in breast and lung cancers. Protein levels for genes of interest were compared in normal (BLUE) versus tumor tissue (RED) using data from UACLAN Portal in breast and lung cancer for **(A)** LOXL2, for COL11A1, COL10A1, COL1A and ADAMTS12 in **(B)** breast cancer and **(C)** lung cancers. Graph represents standard deviations from the median across samples. p values are as indicated and calculated using the t test.

### Pan-cancer analysis of CTHRC1

CTHRC1 (Collagen triple helix repeat containing-1) is a matrisome protein that is most upregulated in cancers (***Figure 1F***). CTHRC1 expression evaluated using GEPIA2 portal confirmed its overexpression in bladder, breast, cervical, colon, lymphoid, brain, kidney, lung, liver, bile duct, ovarian, pancreas, esophagus, cervical, rectal, skin, bone & soft tissue, stomach, testicular, thyroid, thymus, uterine, head and neck cancers (***Figure 5A, C***). Survival analysis showed high CTHRC1 levels to be associated with poor survival in bladder, breast, head & neck, kidney, brain, liver, ovarian, rectal, bone & soft tissue, stomach and thyroid cancers (***Figure 5B, C***). DEG analysis of bladder, breast, head & neck, kidney, liver, ovarian, rectal, stomach, bone and tissue cancers was done and the top 5% genes with a fold change greater or lesser than 2 (p<0.05) selected for further analysis (***Figure 5D***). This identified 19 genes that were upregulated in 3 or more cancers (***Figure 5E-F***), with no genes seen to be downregulated. The 19 upregulated genes include COL11A1, SFRP2, POSTN, EPYC, COMP, COL10A1, OMD, LRRC15, SFRP4, PPAPDC1A, ADAMTS16, OGN, C5orf46, FAP, FNDC1, ODZ3, MMP13, ITGBL1 and THBS4 (***Figure 5F***). They were used to construct a protein protein interaction network using the STRING database (***Figure 5G)***. Functional enrichment analysis of this network revealed ECM organization and adhesion to be among the top 5 biological processes regulated by these network genes (***Figure 5H).*** At the molecular level, these proteins bind Fibronectin, Collagen, cell adhesion molecules and integrins among others (***Figure 5I**)*. Proteins in this network almost exclusively belong to the ECM compartment (***Figure 5J**)*. KEGG analysis of this network further shows their involvement in ECM-receptor interaction, Wnt and focal adhesion signaling (***Figure 5J**)*. Using cytoHubba tool we identified 10 hub genes from this network (POSTN, COMP, COL11A1, MMP13, COL10A1, OMD, OGN, SFRP4, SFRP2 and THBS4) (***Figure 5K**)*.

**Figure 5.**
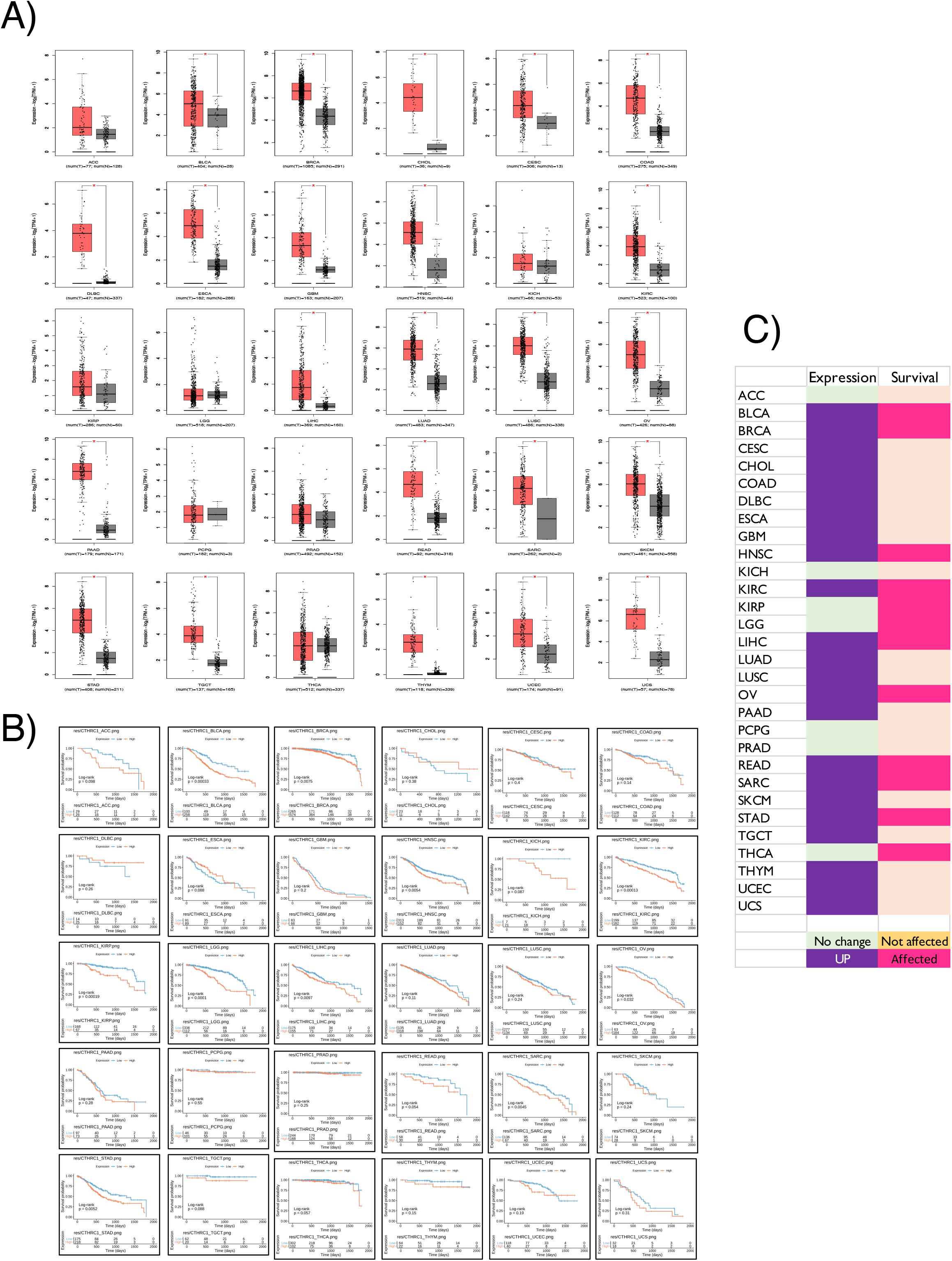
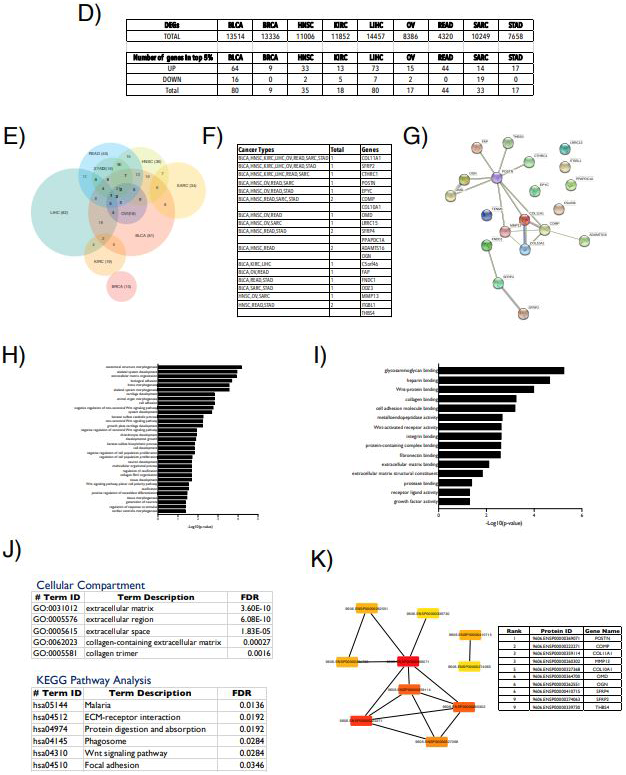
Pan-Cancer analysis of CTHRC1 and differentially expressed genes. **A)** Graphs show expression of CTHRC1 in 30 different cancer types from GEPIA2 database (GREY – normal, RED – tumor). Expression data is represented as mean ± standard deviation (S.D) on a log scale using a box plot. p value of < 0.05 was determined as statistically significant. **B)** Graphs show effect of CTHRC1 on survival in 30 different cancer types using R studio. (BLUE – low expression, RED – high expression). p value of < 0.05 was determined as statistically significant. **C)** Table shows LOXL2 expression and survival data in 30 cancers. Upregulated expression is marked in PURPLE, downregulated expression in BLUE and No change in expression marked in GREEN. Effect seen on survival is marked in PINK and lack of effect marked in CANTALOUPE. **D)** Table lists the TOTAL number of Differential expressed Genes (DEG) in 9 selected cancers (BLCA, BRCA, HNSC, KIRC, LIHC, OV, READ, SARC and STAD – as detailed in Methods). The lower table lists the number of genes in the top 5% of with a fold change of >2 (as detailed in methods) that are differentially up regulated (UP) or downregulated (DOWN). Their aggregate numbers are listed as TOTAL. **E)** This Venn diagram shows the overlap (if any) of these differentially expressed genes in the above listed 9 cancers. **F)** Table lists the 19 overlapping genes between 3 or more cancer types. **G)** Protein protein interaction network constructed for CTHRC1 and its 19 differentially expressed genes using the STRING database shows interactions between CTHRC1 and POSTN and also among several of the differentially expressed genes. BLUE line marks predicted interactions from gene cooccurrence data, GREEN line marks predicted interactions based on gene neighborhood evidence, PURPLE line marks experimentally determined known interactions, YELLOW line marks interactions based on text mining and the LIGHT BLUE line marks interactions based on database evidence. **H)** Gene ontology terms enriched for biological process from STRING network analysis are listed in their descending order of significance. **I)** Gene ontology terms enriched for molecular functions from STRING network analysis are listed in their descending order of significance. **J)** Table lists the gene ontology terms enriched for cellular compartments. Lower table lists the gene ontology terms enriched for protein digestion and absorption using KEGG pathway analysis. The False Discovery Rate (FDR) values from this analysis are listed for each gene ontology term. **K)** Network of 10 hub genes identified using CytoHubba plugin in Cytoscape software. Shown are their protein identifiers from the CytoHubba tool. Colors of the hub genes are based on their rank. These genes are listed as a table in the order of their ranking, high to low.

To validate the significance of CTHRC1 and its hub genes, we first performed a pan-cancer survival analysis of the 10 hub gens along with CTHRC1 using GEPIA2. Of these genes, overexpression of COL11A1, CTHRC1, MMP13, COLA10A1, POSTN and SFRP4, like CTHRC1, is associated with poor prognosis as is low levels of THBS4 (***Figure 6A**)*. We used GEPIA2 to determine if the change in expression of hub genes and CTHRC1 is associated with tumor grade. Statistical analysis of this change by ANOVA showed all hub genes and CTHRC1 expression to indeed be tumor stage dependent (***Figure 6B**)*. Spearman correlation analysis showed a positive correlation between the 10 hub genes and CTHRC1 (***Figure 6C**)*. cBiportal analysis also indicated a co-occurrence between CTHRC1 and OMD, POSTN, OGN, MMP13, COMP and SFRP4 (***Figure 6D**)*. Combining the survival, tumor grade, cooccurrence and correlation analysis, we identified MMP13, SFRP4 and POSTN to be the most likely mediators of CTHRC1 dependent function in cancers (***Figure 6E**)*. Analysis of the UACLAN portal data for breast and lung cancer revealed CTHRC1 protein levels to be upregulated in both cancers (***Figure 7A**)*. In breast cancer samples CTHRC1 was overexpressed with POSTN and MMP13 (***Figure 7B**)*, while in lung cancers POSTN and SFRP4 were overexpressed relative to normal tissues (***Figure 7C**)*.

**Figure 6.**
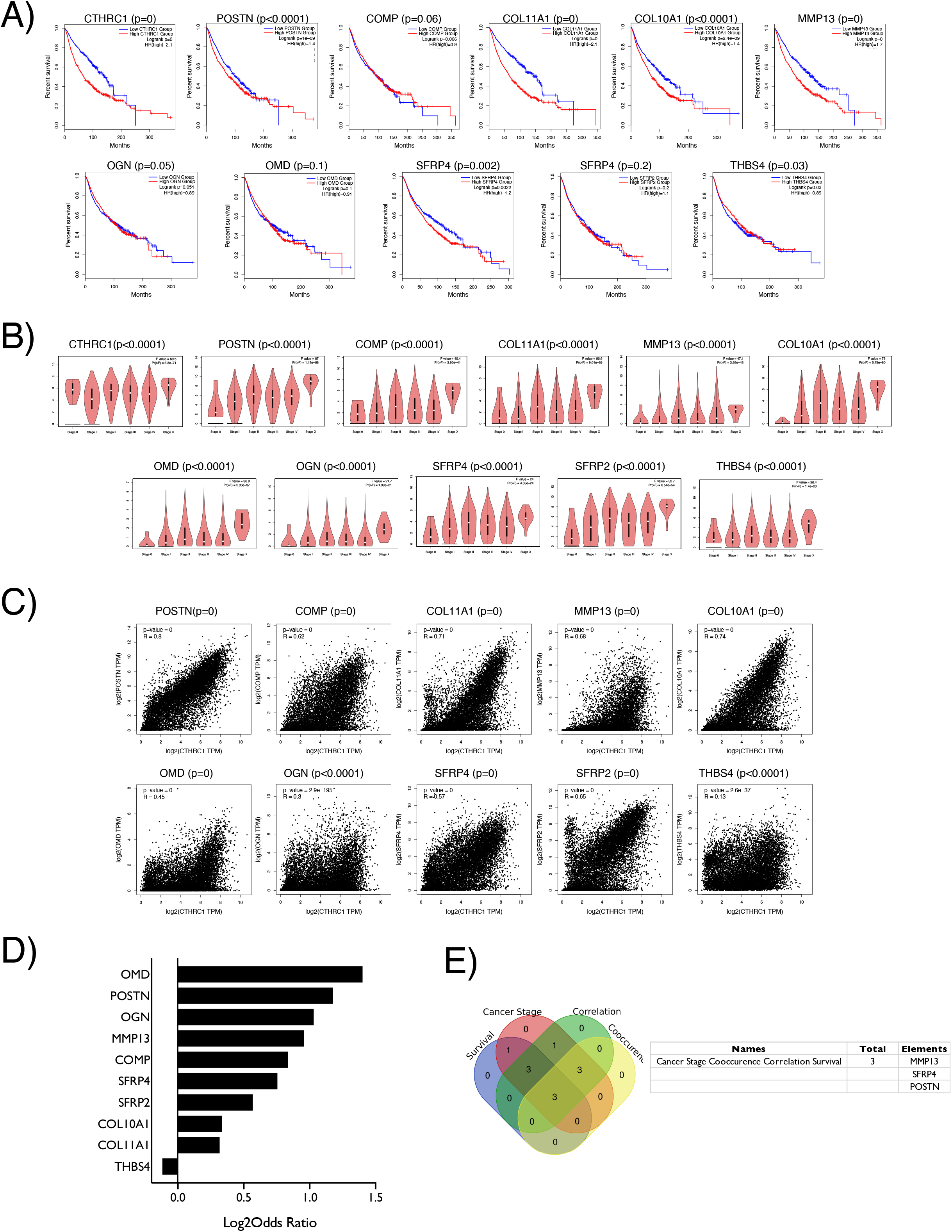
Validation of CTHRC1 and its hub genes in cancers. **A)** Graph represents percentage survival of CTHRC1 and its 10 hub genes (POSTN, COMP, COL11A1, MMP13, COL10A1, OMD, OGN, SFRP4, SFRP2 and THBS2) in 30 cancers using GEPIA2 database. Graph in Blue and Red show the percentage survival over time for cancers with low and high expression respectively of the gene of interest. p values are as indicated above the graph. p values = 0 are representative of very high significance. **J)** Violin plot shows the expression of CTHRC1 and each of its 10 interactors across pathological stages in 30 cancers analyzed using GEPIA2. Significance comparing the stages of cancer for each gene of interest was calculated by ANOVA and significance reported. p values are as indicated above the graph. **K)** Scatter plots show the spearman correlation analysis for CTHRC1 and its interactors in 30 cancers using GEPIA2. p values are as indicated above the graph. p values = 0 are representative of very high significance. **L)** Bar graphs shows log2 odds ratio from cBioPortal for CTHRC1 determining its cooccurrence or mutual exclusivity with the 10 identified genes of interest (POSTN, COMP, COL11A1, MMP13, COL10A1, OMD, OGN, SFRP4, SFRP2 and THBS2) in 30 cancer types. **M)** This Venn diagram shows the overlap between genes that in a statistically significant manner affect survival, affect cancer staging, are related in correlation analysis and show cooccurrence in 30 cancer types. Table lists the 3 genes overlapping in all four analysis.

**Figure 7.**
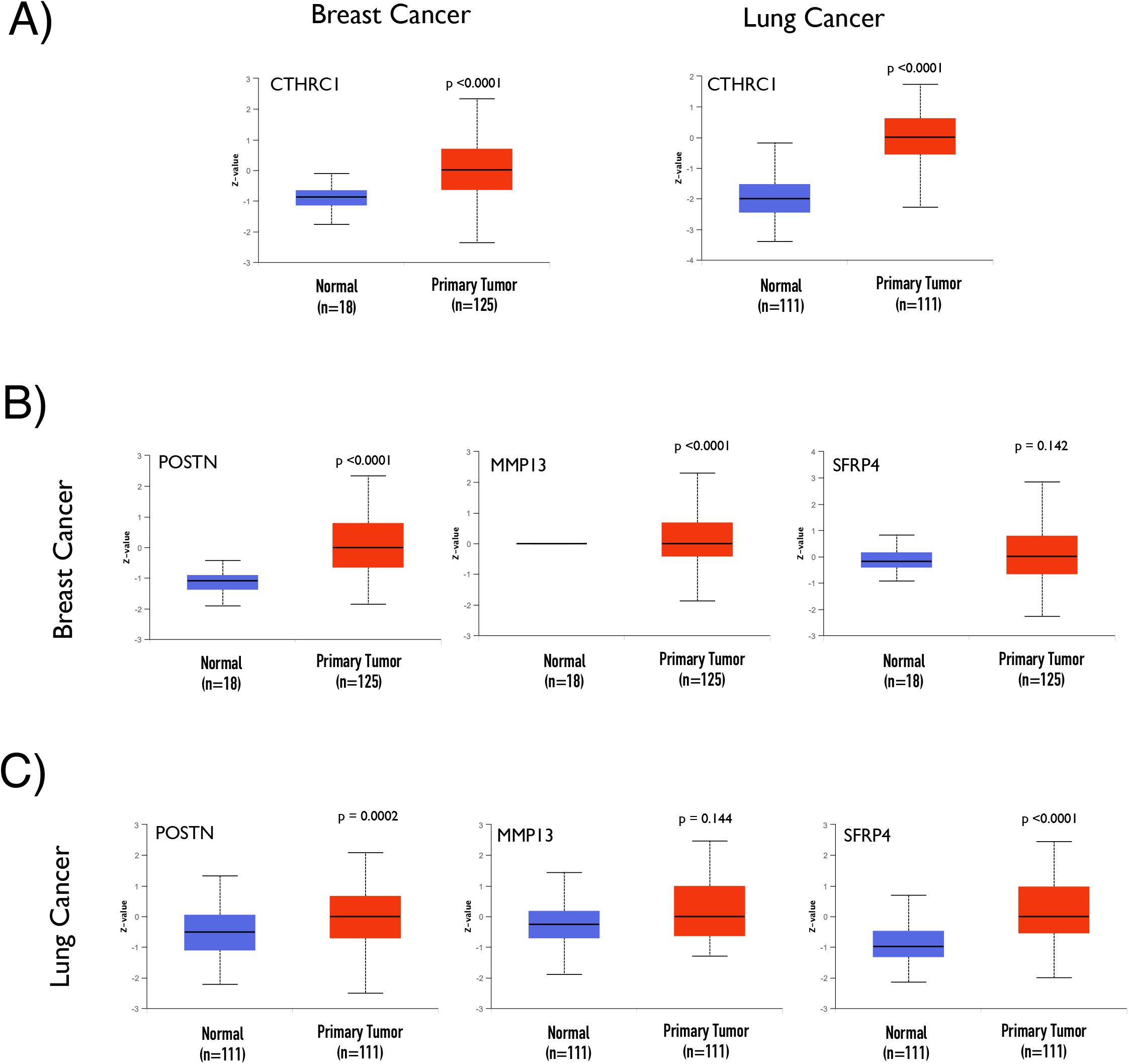
Protein levels of CTHRC1, POSTN, MMP13 and SFRP4 in breast and lung cancers. Protein levels for genes of interest were compared in normal (BLUE) versus tumor tissue (RED) using data from UACLAN Portal in breast and lung cancer for **(A)** CTHRC1, for POSTN, MMP13 and SFRP4 in **(B)** breast cancer and **(C)** lung cancers. Graph represents standard deviations from the median across samples. p values are as indicated and calculated using the t test.

Together these observations identify CTHRC1 along with POSTN, SFRP4 and MMP13 along with LOXL2 and its interactors COL11A1, COL10A1, COL1A1 and ADAMTS12 to be vital mediators of the impact the tumor ECM has across cancers.

## Discussion

In the past decade the number of publications on ECM and cancer has doubled and ECM has emerged as a key player in progression, diagnosis and treatment of cancers. The tumor ECM is primarily secreted by the cancer associated fibroblasts (CAF) and supports the formation of a premetastatic niche [24,25]. Changes in the ECM composition and structure (i.e., reorientation of collagen fibrils) have been known to promote migration and invasion of cancer cells [24,26] [2][21]. Altered ECM deposition is further associated with poor prognosis in multiple cancers [2,3]. Our pan-cancer study, using TCGA data from 30 cancer types, takes a stringent look at multiple aspects of the regulation of 1027 matrisome genes, including their copy number, gene expression and effect on survival. The top 5% of genes with the most amplifications or deletions, which show altered expression in 50% or more cancers and affect survival in 20% or more cancers, were selected. This identified CTHRC1, LOXL2, SNED1, IGFBP3, BMP1, PI3, COL1A2 and AEBP1 as the top 8 pan-cancer matrisome genes of interest. Of these LOXL2 and CTHRC1 were the most prominently upregulated and SNED1 the most prominently downregulated gene across cancers. Our intent was to look at the regulatory network of interactors for these matrisome proteins that together could mediate the effect ECM has across cancers.

SNED1 was recently discovered as a ECM glycoprotein whose exact function remains unknown [22,27]. SNED1 knockout mice die perinatally showing significant growth defects [22]. SNED1 is highly expressed by mammary tumors in mice, its knockdown causing fewer lung lesions [4], though the mechanisms by which it regulates cancer progression remains largely unknown [4]. We have identified SNED1 to be downregulated and its low levels is associated with poor survival in ~36% of cancers. Our stringent DEG analysis to identify genes that could be potentially regulated by SNED1 in three or more cancers did not yield any candidates of interest. *In silico* studies predicting possible binding partners of SNED1 have suggested the presence of an RGD and LDV motif in SNED1 could potentially support integrin binding [27]. Interactome studies predict multiple forms of collagen to also be possible binding partners for SNED1 [27], raising the possibility that it could be involved in regulating the assembly and organization of collagen.

CTHRC1 and LOXL2 are both also known regulators of collagen synthesis and assembly [23,28]. Collagen makes up at least 30% of the total protein in the human body [29] and is highly aligned and anisotropic contributing to tumor cell survival and metastasis [30]. Changes in collagen crosslinking and architecture promote malignancy and cancer cell dissemination [30,31]. CTHRC1 has been shown to inhibit collagen type I and III transcripts [32], with KO mice showing reduced type I collagen levels [33]. LOXL2 is also a major regulator of collagen, affecting fibril alignment, directionality, orientation and width [30]. High levels of LOXL2 in cancers are associated with increased metastatic potential possibly due to their effect on tumor cell invasion and migration [61]. In breast cancer, increased LOXL2 expression by the tumor stroma is associated with poor prognosis [34]. Most published studies have focused on the canonical enzymatic function of LOXL2 in regulating collagen crosslinking and fibril formation, supporting tumorigenesis [23]. Further supporting this role for LOXL2 our study identifies COL11A1, COL10A1, COL1A1, ADAMTS12 and PPAPDC1A as possible interactors of significance across cancer types.

COL11A1 highly expressed by CAFs is upregulated in breast, esophageal, gastric, lung, ovarian and colorectal cancers and also associated with poor survival, drug resistance and cancer recurrence [35–37]. High COL10A1 levels are reported in multiple solid tumors including breast, colon, bladder, stomach, esophagus, lung, testis, ovary and pancreas[38] and associated with poor prognosis in breast and lung cancers [39,40]. COL1A1 overexpression is similarly reported in breast and lung cancer and associated with metastasis and poor survival [41,42]. ADAMTS12, a member of the disintegrin and metalloproteinase with thrombospondin repeats gene family, has been shown to have dual roles both as tumor promoter and tumor suppressor [43]. Members of the ADAMTS family of proteolytic enzymes are implicated in a variety of physiological processes, including collagen maturation [44,45]. They also play significant role in the progression of cardiovascular diseases, autoimmune disease and cancer [43–46]. PPAPDC1A or PLPP4 (phospholipid phosphatase) is an integral membrane glycoprotein which is speculated to be involved in regulating tumor progression [47]. PLPP4 has been shown to be highly expressed in lung cancer and its knockdown in cells inhibits cell proliferation. High PLPP4 expression is associated with poor prognosis in lung cancer patients [48].

Of these possible interactors of LOXL2, COL1A1 levels are seen to be regulated by LOXL2 in normal mouse fibroblasts [49,50]. COL11A1, COL10A1 and COL1A1 have been shown to be a part of tumor matrisome signature genes that are consistently upregulated in multiple cancers [14,15,51]. Studies looking at role of COL11A1 in cancers have identified COL10A1, COL1A1, LOXL2 and ADAMTS12 expression to be correlated with COL11A1 [52]. Together these observations further emphasize the joint role these genes could have across cancers. Besides the direct regulation by LOXL2 of collagen (COL11A1, COL10A1 and COL1A1), the possible role of ADAMTS12 and PPAPDC1A working in tandem with LOXL2 across cancers, needs to be explored. Recent studies have shown that LOXL2 also has intracellular functions independent of its enzymatic activity. LOXL2 is seen to inhibit EMT possibly by binding and repressing E-cadherin promoter in HEK293T cells [53]. The impact of a non-canonical role for LOXL2 in regulating collagen synthesis, assembly and function also remains to be investigated.

CTHRC1 is similarly seen to work with possible interactors POSTN, MMP13, SFRP4 across cancers. Studies have shown CTHRC1 to be overexpressed in lung [54], breast [55], skin [56], gastric [57], liver [58], endometrial [59] cancers and in glioma [60], and its expression associated with advanced disease stage and decreased survival. POSTN an ECM protein is highly expressed by the tumor stroma and associated with poor prognosis in breast and lung cancer, affecting survival rates and metastasis [11]. Though CTHRC1 and POSTN upregulation in breast cancer is associated with poor prognosis [55], no direct interaction or correlation between these genes has so far been reported. MMP13 (collagenase 3) known to act on collagens (collagen type I, II, IV and IX) contributes to increased angiogenic potential in tumors [61,62] and its high expression correlates with poor prognosis across several cancers [63]. In breast and lung cancers, high levels of MMP13 are associated with metastasis [63,64]. SFRP4 is highly expressed by the tumor stroma in various cancers and in breast cancer its differential expression across tumor stages is indicative of poor survival [65].

POSTN and MMP13 are known to independently regulate collagen, but these genes are not reported as mediators of CTHRC1. Network analysis in this study suggests their possible crosstalk emphasizing the need to explore their joint role across cancers. Similarly, SFRP4 (a known Wnt antagonist) could also regulate collagen architecture and assembly by controlling Wnt signaling via beta catenin [66]. CTHRC1 is a known regulator of the Wnt pathway during development [67]. Changes in Wnt signaling are known to drive the aggressiveness of breast and lung cancers [68]. These observations when taken in context of this study support a joint role for CTHRC1, POSTN, MMP13, SFRP4 in collagen organization possibly dependent on Wnt signaling.

Collagens are the most abundant protein in the human body accounting for as much as 90% of connective tissue [29]. Synthesis of collagen involves a number of enzymatic post-translational modifications [69,70].Collagen fibrils are further strengthened by the covalent crosslinking between lysine residues by lysyl oxidases (LOX) [70,71]. This study in carefully examining the role of the matrisome across cancers, has identified CTHRC1 and LOXL2 as major ECM regulators across cancers. Both play a role in collagen synthesis and assembly [23,28] by possibly interacting with POSTN, MMP13, SFRP4, COL11A1, COL10A1, COL1A1, ADAMTS12 and PPAPDCA1. This suggests the regulation of collagen could be a vital pan-cancer mediator of disease progression. The extensive cross-linking of collagen fibers regulates the density, packing order of collagen which in turn could contribute to changes in ECM stiffness that may further affect cancer progression [72–77]. CTHRC1 and LOXL2 are both also secreted and detectable in the blood [10,78] making them potential biomarkers of interest across cancers.

## Acknowledgements

We would like to thank Dr. Mikhail Dozmorov (Virgnia Commonwealth University) for use of his code from the GitHub repository and also for helping us troubleshoot the code.

